# REM Sleep Movements in Parkinson’s Disease: Is the Basal Ganglia Engaged?

**DOI:** 10.1101/2022.05.25.493494

**Authors:** Ajay K. Verma, Sergio Francisco Acosta Lenis, Joshua E. Aman, David Escobar Sanabria, Jing Wang, Amy Pearson, Meghan Hill, Remi Patriat, Lauren E. Schrock, Scott E. Cooper, Michael C. Park, Noam Harel, Michael J. Howell, Colum D. MacKinnon, Jerrold L. Vitek, Luke A. Johnson

## Abstract

To elucidate the role of the basal ganglia during REM sleep movements in Parkinson’s disease (PD) we recorded pallidal neural activity from four PD patients. Unlike desynchronization commonly observed during wakeful movements, beta oscillations (13-35 Hz) *synchronized* during REM sleep movements; furthermore, high-frequency oscillations (150-350 Hz) synchronized during movement irrespective of sleep-wake states. Our results demonstrate differential engagement of the basal ganglia during REM sleep and awake movements.

---

The basal ganglia play a critical role in the control of motor function during wakefulness^1,2^. Voluntary movements are preceded by desynchronization (reducing in power) of beta oscillations (13-35 Hz) in the globus pallidus internus (GPi)^3,4^. Excessive spontaneous synchronization of beta oscillations (increasing in power) in the GPi in people with Parkinson’s disease (PD) is thought to contribute to akinesia and bradykinesia since the suppression of beta power by dopamine replacement therapy or deep brain stimulation (DBS) improves motor behavior^4–6^. High-frequency oscillations (HFOs, >100 Hz) in the GPi have been observed to synchronize during wakeful movement in unmedicated PD patients^4,7^. Furthermore, a recent study in PD patients on and off medication, and in naïve and parkinsonian non-human primates suggests that exaggerated HFOs have pathophysiological relevance in PD^4^. Our understanding regarding the modulation of beta oscillations in the basal ganglia during REM sleep movements is limited^8^, however, and HFOs in such context have not been investigated.

In contrast to wakefulness, people with PD can demonstrate improved motor activity during rapid eye movement (REM) sleep^9^. In particular, a high percentage of individuals with PD have REM sleep behavior disorder (RBD), a parasomnia characterized by the loss of muscle atonia and a dramatic increase in motor activity, often with punching and kicking behavior suggestive of dream enactment^10^. People with PD without a clinical diagnosis of RBD can also demonstrate motor activity during REM sleep^9,11^. Currently, little is known about the role of the basal ganglia in the control of movements that occur during REM sleep. Recordings from the subthalamic nucleus have provided initial evidence of marked differences in the dynamics of movement-related oscillations between wake and sleep states^8^. To further explore the role of the basal ganglia in movement control during REM sleep, we recorded local field potentials (LFPs) from four PD patients via externalized directional deep brain stimulation (DBS) leads implanted in the GPi (the principal output nucleus of the basal ganglia). We characterized the dynamics of beta and HFOs in the GPi during REM sleep and awake movements in an effort to improve our understanding of the functional role of the basal ganglia oscillations underlying movement across sleep-wake states.

All subjects were noted to display body movements during REM sleep as identified by video-polysomnography. The number of movements identified during REM sleep, predominant motor phenomena observed, and other demographic information for each subject are reported in **Table 1**. One subject (#2) had a documented history of sleep dysfunction and displayed complex movements suggestive of dream enactment.

**Table 1.**
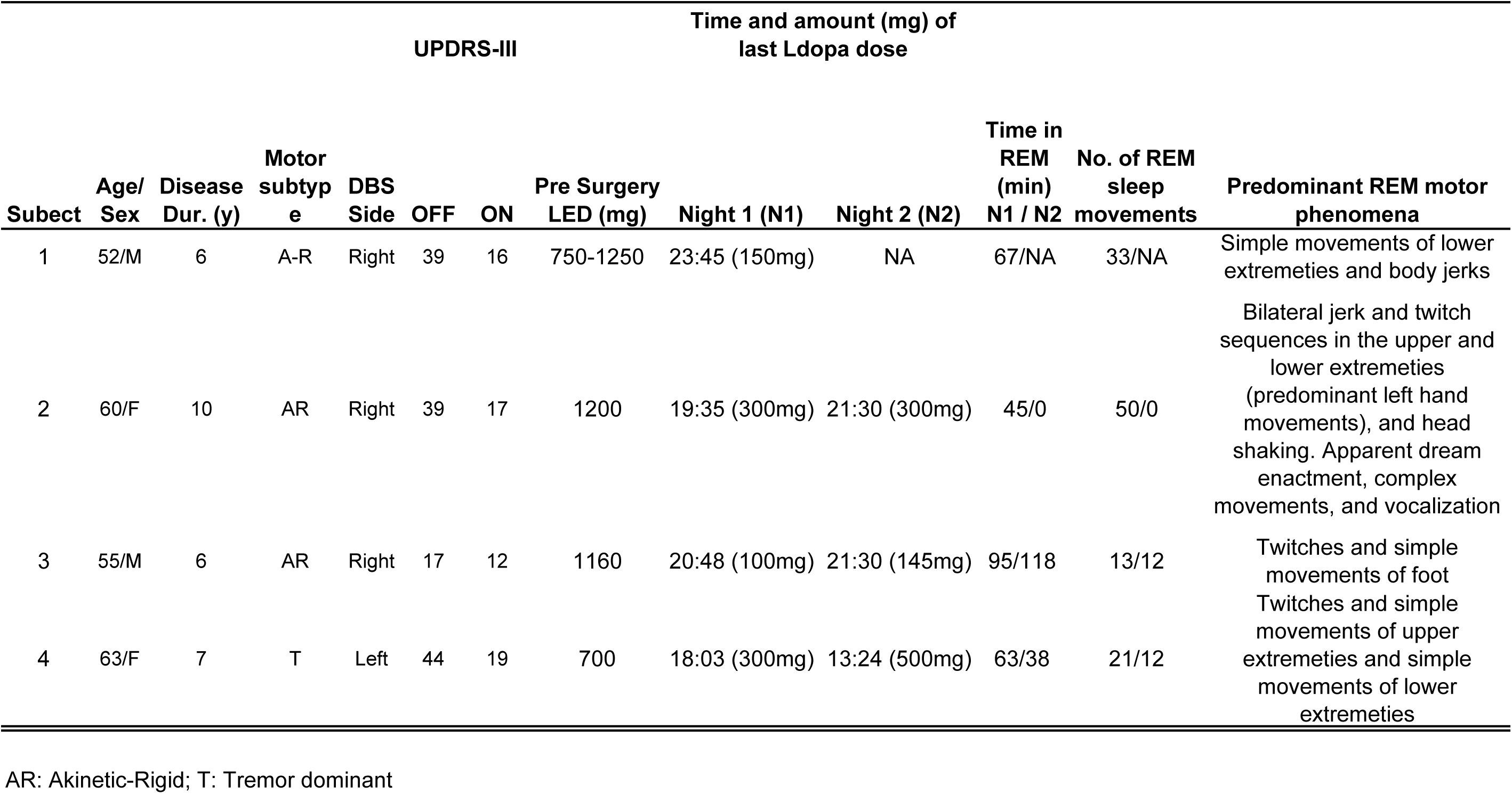
Demographic data at time of study.

The contact pair on the directional DBS lead exhibiting the highest modulation of beta oscillations for each patient during awake voluntary movements was chosen for analysis and reported in **Figure 1**. Additional details regarding beta and HFO modulation associated with REM sleep and awake movements for other contact pairs on the directional DBS lead, and lead location in the GPi, are summarized in **Supplemental Figure 1**.

**Figure 1.**
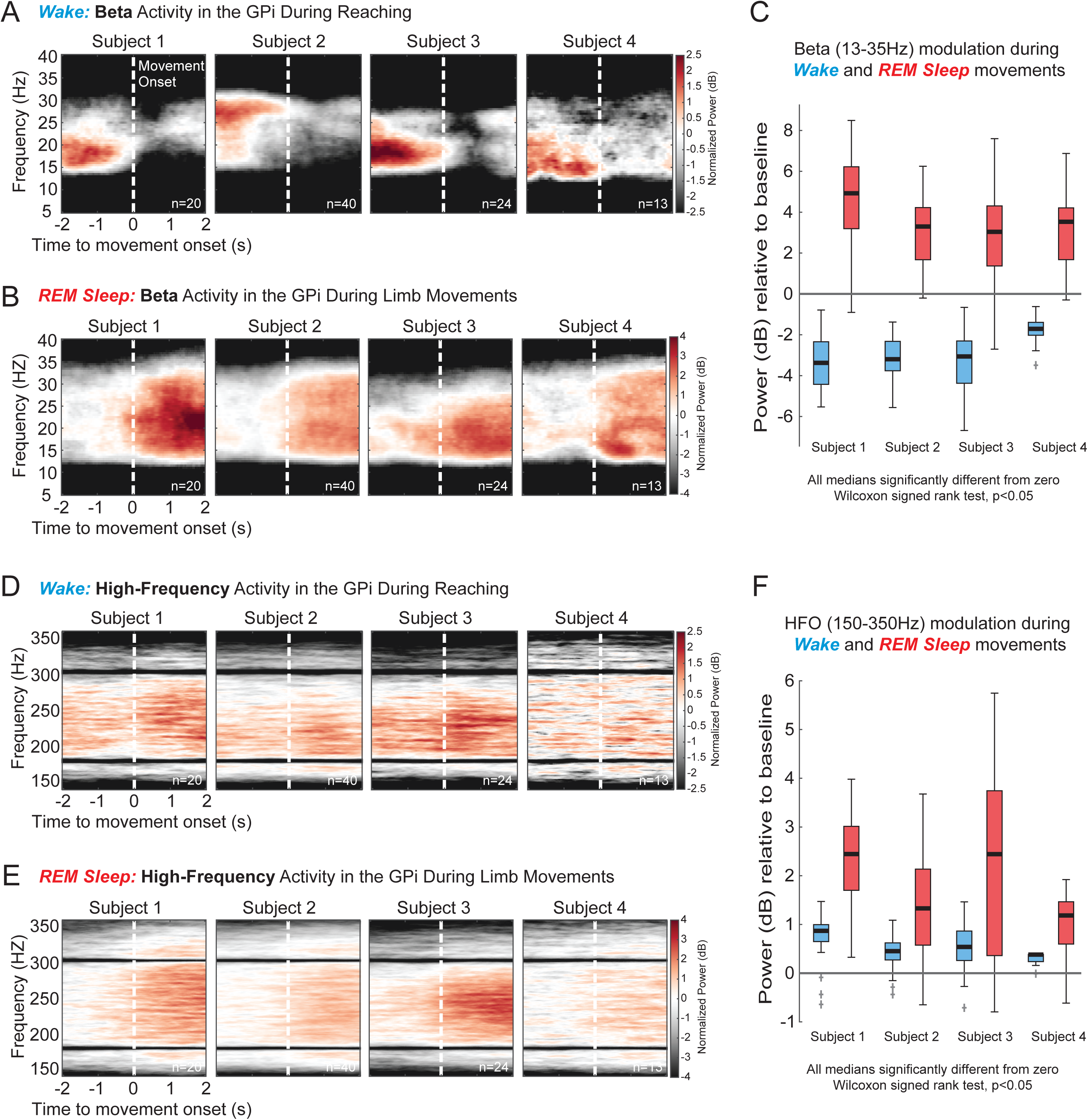
Movement-related beta and high-frequency oscillations recorded from DBS leads in the GPi of PD patients in awake and REM sleep states. **(A)** Trial-averaged spectrograms aligned to movement onset showing beta (13-35 Hz) desynchronization in the GPi during wakeful volitional movement (reaching task, see Supplemental Methods). **(B)** Spectrograms showing beta synchronization in the GPi during REM sleep movements. **(C)** Distributions of beta band power modulation (relative to pre movement baseline) during wake movements (blue) and REM sleep movements (red). All data distributions were significantly different from zero (Wilcoxon signed rank (WSR) test, p<0.05). **(D, E)** Trial-averaged spectrograms aligned to movement onset show synchronization of high-frequency oscillations (HFO, 150-350Hz) in the GPi during wakeful and REM sleep movements, respectively. **(F)** Distributions of HFO band power modulation (relative to pre movement baseline) during wake movements (blue) and REM sleep movements (red). All data distributions were significantly different from zero (WSR test, p<0.05).

Trial-averaged spectrograms depicting oscillatory power changes in the beta band (13-35 Hz) during awake voluntary movements (reaching task, see Supplemental Methods for details) and REM sleep movements are shown in **Figure 1A** and **1B**, respectively. In the awake condition, beta oscillations significantly desynchronized during movement execution in all subjects compared to a pre-movement period (Wilcoxon signed rank (WSR) test, p<0.05, **Figure 1C**, blue boxplots). Conversely, all participants exhibited strong synchronization of beta oscillations during REM sleep movements compared to baseline (WSR test, p<0.05 **Figure 1C**, red boxplots). In contrast to the beta band, which showed opposite polarity of modulation during awake and REM sleep movements, HFOs (150-350 Hz) showed movement-related synchronization during both awake voluntary and REM sleep movements (**Figure 1D, 1E**).

The primary finding of this study was that, in people with PD, movement-related beta oscillations in the GPi synchronized during REM sleep but desynchronized during wakefulness. In contrast, HFOs in the GPi synchronized during movement regardless of sleep-wake state. Taken together, our results demonstrate that GPi is engaged during REM sleep movements and question the hypothesis that the basal ganglia is not involved in the generation of movements during REM sleep in people with PD^9^.

Previous studies have reported improved motor behavior (e.g. faster, smoother movements) during REM sleep in PD patients with RBD and have hypothesized a model where REM sleep movements are generated by the extrapyramidal pathways bypassing the dopamine deficient basal ganglia^9,12^. Potential support for this idea comes from a study in four PD patients with RBD showing movement-related synchronization of the subthalamic nucleus (STN) beta oscillations during REM sleep^8^. Since STN beta oscillations in the basal ganglia desynchronize during wakeful movements, which is thought to be permissive to the movement^3,8,13^, the authors interpreted their data as supportive of the hypothesis that pathological basal ganglia signaling is bypassed during REM sleep and that the movements during REM sleep are mediated by pathways alternate to the basal ganglia-cortical motor network in people with PD. We also observed significant synchronization of beta oscillations in the GPi during REM sleep movements in all four subjects, which could be viewed as providing additional support for this hypothesis. Our findings of HFO synchronization in the GPi in both awake and sleep conditions, however, complicate this interpretation. Elevated HFO power has been shown to be a characteristic feature of movements during wakefulness in PD^4,7,14^. If movements during REM sleep are not mediated by the basal ganglia, then the polarity of modulation of HFOs during REM sleep movement might be expected to be opposite to what was observed during wakefulness as was observed with beta oscillations. However, the polarity of movement-related modulation of HFOs in the GPi during REM sleep was similar to that during wakefulness i.e., synchronizing during movement irrespective of sleep-wake states. We interpret these findings as demonstrating that the basal ganglia is likely engaged during REM sleep movements, but its role in movement control differs greatly during wakefulness and REM sleep.

Furthermore, our observation that polarity of modulation in beta and HFOs between PD patients with (n=1) and without (n=3) RBD were qualitatively similar leads us to speculate that synchronization of beta oscillations and HFOs in response to movements generated during REM sleep are characteristic of REM sleep in PD and may not be unique to people with PD diagnosed with RBD. A future study with a higher sample size of PD patients with and without RBD will be required to confirm this observation.

Despite our understanding of skeletal muscle atonia during REM sleep, small movements during REM sleep is common in healthy individuals^15,16^. Whether the movement-related changes in beta oscillations and HFOs we observed during REM sleep are specific to PD or more generally applicable remains unclear. Identification of these events with concomitant recordings across the basal ganglia-cortical motor network can shed further light on the neural mechanisms of movement control during REM sleep and enhance our understanding of physiological and pathological movements generated during sleep. While it is not feasible to perform such electrophysiological recordings in healthy controls, data collection may be feasible from patients with non-PD neurological disorders such as dystonia and Tourette syndrome, where the STN and GPi are common DBS targets^5,17–19^, to understand if movement-related dynamics of beta oscillations and HFOs during REM sleep is unique to PD.

In summary, we showed that, contrary to the stereotypical pattern of movement-related beta desynchronization in the GPi during voluntary wakeful movements, beta oscillations in the GPi are synchronized during REM sleep movements. Furthermore, HFOs are synchronized during movement irrespective of sleep-wake state. Together, our results demonstrate differential engagement of pallidum during wake and REM sleep movements and these findings may inform the development of DBS approaches tailored to suppress excessive movements generated during REM sleep in people with PD.

## Methods

This study was approved by the University of Minnesota Institutional Review Board (#1701M04144) and informed consent was obtained according to the Declaration of Helsinki. Four participants (two female) with idiopathic PD consented to externalization of their DBS lead. All subjects were implanted unilaterally with a directional DBS lead targeting the GPi. Details of the surgical procedures for GPi DBS implantation and externalization are described in the Supplemental Methods. LFPs were recorded over the course of a 48-hour period (4-8 days after lead implantation) from the externalized GPi lead along with video-polysomnography (PSG), which included video, electrooculography (EOG) of both eyes, chin electromyography (EMG), and electroencephalography (EEG) according to 10-20 electrode placement. Sleep stages and ancillary events were scored according to the American Academy of Sleep Medicine Scoring of Sleep and Associated Events Version 2.6^20^. Detailed descriptions of data collection and analysis are provided in the Supplemental Methods.

## Acknowledgments

We thank the participants in this study for their time and willingness to contribute to this research. We thank the entire University of Minnesota Udall team for their support of this study. We would like to acknowledge: Eric Maurer for leadership in Udall project management; Kelly Brown and Kelly Ryberg for patient recruitment and project management; Greg Molnar for his role in supporting initial investigations of sleep using DBS lead recordings; Ethan Marshall, Sinta Fergus, Stephanie Alberico, and Kevin O’Neill for help during data collections. This work was supported by the Udall Center for Excellence in Parkinson’s Disease, National Institutes of Health, National Institute of Neurological Disorders and Stroke: P50-NS123109, P50-NS098573, R01-NS110613, R01-NS058945, R01-NS037019, P30-NS076408, P41-EB027061; University of Minnesota’s NIH Clinical and Translational Science Award: UL1TR002494; MnDRIVE (Minnesota’s Discovery, Research and Innovation Economy) Brain Conditions Program; Engdahl Family Foundation; The Kurt B. Seydow Dystonia Foundation.

## Authors Contribution

1. Research project: A. Conception, B. Organization, C. Execution;
2. Analysis: A. Design, B. Execution, C. Review and Critique;
3. Manuscript Preparation: A. Writing of the first draft, B. Review and Critique;

AKV: 2A, 2B, 2C, 3A, 3B

SAL: 2A, 2B, 2C, 3A, 3B

JEA: 1B, 1C, 3B

DES: 1B, 1C, 3B

JW: 1B, 1C, 3B

AP: 2B, 3B

MH: 1C, 3B

RP: 1C, 2A, 2B, 2C, 3B

LES: 1C, 2C, 3B

SEC: 1C, 2C, 3B

MCP: 1A, 1C, 2C, 3B

NH: 1A, 1C, 2A, 2B, 2C, 3B

MJH: 2A, 2B, 2C, 3A, 3B

CDM: 1A, 2C, 3A, 3B

JLV: 1A, 1B, 1C, 3B

LAJ: 1A, 1B, 1C, 2A, 2B, 2C, 3A, 3B

## Data Availability Statement

The data that support the findings of this study are available from the corresponding author upon reasonable request.

## Figure and Supplementary Figure Legends

**Figure S1. GPi DBS recording locations and movement-related beta and high-frequency power modulation by direction. (A)** Schematic of an Abbott directional “1–3–3–1” lead, illustrating in orange that bipolar paired recordings from vertically adjacent segments were used in this study. **(B)** DBS lead implant locations for each patient, estimated from preoperative MRI and postoperative CT scans. Axial reconstructions are shown, with recording segment used in primary analysis (**Figure 1**, Results section) shown in orange, chosen based on the segment direction with largest movement-related modulation of beta band activity (see Supplemental Methods). Recordings were made in right GPi in patients 1-3 and in left GPi in patient 4; left GPi images were mirrored for visualization. Subject 3 was implanted with a Boston Scientific directional lead (See Supplemental Methods) which has alternative contact labeling compared to Abbott, however for visualization in this figure, directions were assigned A, B and C labels. Distributions (lower panels) and median magnitude (upper panels) of beta band modulation in each direction are presented. To determine statistically significant directionality of oscillatory activity, the Kruskal-Wallis test was performed separately on wake and REM sleep datasets to determine if distributions in the three directions come from the same distribution (p<0.05), followed by pairwise tests correcting for multiple comparisons. Analogous plots but for high-frequency band power modulations are shown in **(C)**.

