## Supplemental Figure 1 for "REM Sleep Movements in Parkinson’s Disease: Is the Basal Ganglia Engaged?"

**Supplemental Material**

**Methods**

**Data collection**

*Surgical Procedure:* Details of the surgical procedures for GPi DBS implantation and lead externalization are described in detail in a previous publication[1,2]. Briefly, subjects underwent standard 3T MRI (all subjects) and a high resolution 7T MRI (excluding Subject 3) for direct targeting and postoperative lead localization[3,4]. Intraoperative electrophysiological mapping techniques were used to identify the sensorimotor region of GPi for implantation[5]. In all subjects a directional “1-3-3-1” electrode was used (Subjects 1,2,4: Abbott Infinity model 6172, illustrated in **Supplemental Figure 1A**; Subject 3: Boston Scientific Vercise Cartesia model DB-2202-45; all leads had 1.5 mm contact height with 0.5 mm vertical spacing). After implantation, the lead was connected to an extension wire which was tunneled to a subcutaneous pocket in the chest and then connected to another extension wire which was externalized at the abdomen (“percutaneous extension”)[1]. Externalized components were secured and protected with a water-proof barrier dressing and subjects were discharged to home to recover. Externalization recordings occurred 4-8 days later, allowing some time for reduction of microlesion effects that can occur following lead placement[6,7]. After the study, subjects returned to the hospital for removal of the percutaneous or externalized extension wire and placement of the IPG. A movement disorders clinician performed DBS programming approximately 4-6 weeks after IPG placement per standard clinical care.

*Sleep Recording and Staging:* Local field potential (LFP) activity from the DBS lead and scalp EEG (10-20 montage, modified as needed to accommodate scalp incision) were collected over the course of two days while residing in the University of Minnesota Health Clinical Research Unit. Signals were recorded on an Xltek NeuroWorks Workstation (Quantum amplifier, 4096 Hz sampling rate, Natus) to enable post-hoc video-polysomnography (v-PSG) and analysis of time-synchronized pallidal oscillatory activity.

v-PSG data (including EEG, EOG, EMG, video) were imported into Natus SleepWorks software. Sleep staging was performed according to the American Academy of Sleep Medicine (AASM) Manual for the Scoring of Sleep and Associated Events Version 2.6 by a registered polysomnographic technician (A.P.) using the standard EEG, EOG, and chin EMG montage[8]. Sleep stages (Wake, NREM 1 (N1), NREM 2 (N2), NREM 3 (N3), and REM) were first determined in standard 30-second epochs (consistent with AASM scoring guidelines). Subsequently, 10-second mini-epochs were subdivided to better identify brief sleep phenomena including K complexes, sleep spindles, slow-wave activity, phasic REM, and tonic REM. Neural data and sleep stages and sleep phenomena annotations were then exported in EDF format for subsequent analysis.

*REM sleep movement and reaching task:* v-PSG recordings were reviewed by experienced sleep specialists (M.H. and A.P.). Movements occurring during REM sleep epochs were visually identified and characterized (twitches, periodic limb movements, complex dream enactment, limb adjustment, or body position change) and timestamps of movement onset were saved for further analysis of event-related potentials. Predominant motor phenomena for each subject are described in Table 1. As a comparison condition, GPi LFP data collected during a daytime motor task were also analyzed. LFP data were collected while patients performed a simple touchscreen reaching task using the arm contralateral to the implanted DBS lead. Trials began with the hand on a digitized home button located 45 cm from a touchscreen monitor. After a randomized variable 3-4 sec delay following the start of a trial, a 1.27 cm hollow circle (target) appeared on the center of the touchscreen along with a 5 cm square box directly to the left of the circle (10 cm). The appearance of the circle and square was the patient's "go cue". Subjects were instructed to touch and drag the circle into the square box as quickly and accurately as possible and then return to the home button. The number of reach trials included in the analysis were matched to the number of REM movements detected in each patient and are reported in the figures (see Data analysis and statistics section below for additional details). Wake task-related data were included in previous publications but analyzed differently in the present study[1,2].

**Data analysis and statistics**

All analyses were performed using customized scripts in MATLAB (MathWorks). LFP activity was extracted via bipolar montage (i.e. signal subtraction) of vertically adjacent DBS contacts within the GPi (see **Supplementary Figure 1A**); the segmented contacts of the 1-3-3-1 directional lead were used in this study, resulting in three bipolar pairs that were analyzed.

A multimodal exploration of video, EEG (including EOG and EMG), and LFP signals (in time and frequency domain) was done to identify artifact-free movement events during REM. Timestamps of movement initiation during the reaching task were similarly identified. In all subjects, the number of reach trials (n=50) surpassed the number of identified REM sleep movements. The same number of artifact-free samples were collected in chronological order from the data associated with the reaching task (i.e. if 40 REM sleep movements were detected, the first 40 reach movements were analyzed).

LFP signals were filtered between 13-35 Hz for the beta oscillations, and between 150-350 Hz for the high-frequency oscillations (HFOs). Perievent spectrograms aligned to movement onset were computed via the multi-taper method and Chronux toolbox using a moving window of 1s, 10ms steps, with three tapers resulting in a frequency resolution of 0.5 Hz[9]. For each subject and frequency band (i.e. beta, HFO) the spectrogram was normalized to the total power in the target frequency band in a 1 sec baseline period and reported in units of dB relative to the baseline[10,11]. To quantify movement-related changes in the normalized spectrogram, for each trial the mean band power in a pre-movement period 2 seconds before movement onset ($AvgPower_{Pre\text{-}move}$) and in a movement period 2-sec duration beginning with movement onset ($AvgPower_{Move}$) were calculated. Boxplots display distributions of normalized trial-by-trial movement-related band power modulation ($AvgPower_{Move} - AvgPower_{Pre\text{-}move}$). Non-parametric statistical tests were performed because of the relatively small sample size and the non-normality of the data. To determine whether a subject's median movement-related modulation was significantly different from zero, reflecting significant synchronization (positive value) or desynchronization (negative value) in the frequency band of interest, the Wilcoxon signed-rank test was performed ($p < 0.05$). The segment pair in each subject with the greatest movement-related modulation in the beta band is presented in the main results section and **Figure 1**. Movement-related modulation from all three segment pairs for each subject is presented in **Supplementary Figure 1**. To test whether there is some spatial specificity between recording directions in the GPi, the sleep and wake movement power modulations from the three directions were compared using the Kruskal-Wallis test ($p < 0.05$), testing the null hypothesis that all three distributions come from the same distribution, with post-hoc pairwise comparisons between directions made with Bonferroni correction for multiple comparisons.

**Determining DBS lead orientation**

DBS lead locations in the GPi were estimated based on information obtained during intraoperative electrophysiological mapping as well as co-registered preoperative MRI and postoperative CT scans (see reference # [4] for details). The orientation of the DBS lead and relative direction of individual segments for each patient were derived from the fiducial marker on the lead, in combination with the unique artifact characteristics of the segments, using a modified version of the DiODe algorithm[12]. The original DiODe algorithm was designed and validated for the Boston Scientific Cartesia electrodes; since most of our patients were implanted with the Abbott Infinity electrodes, the MATLAB code was modified to be compatible with the new electrode characteristics (e.g., smaller marker and slight changes in the intensity profiles of the segments artifacts) in collaboration with Dr. Dembek, the lead author of the DiODe algorithm (personal communications, R.P.). Additional details can be found in[2].

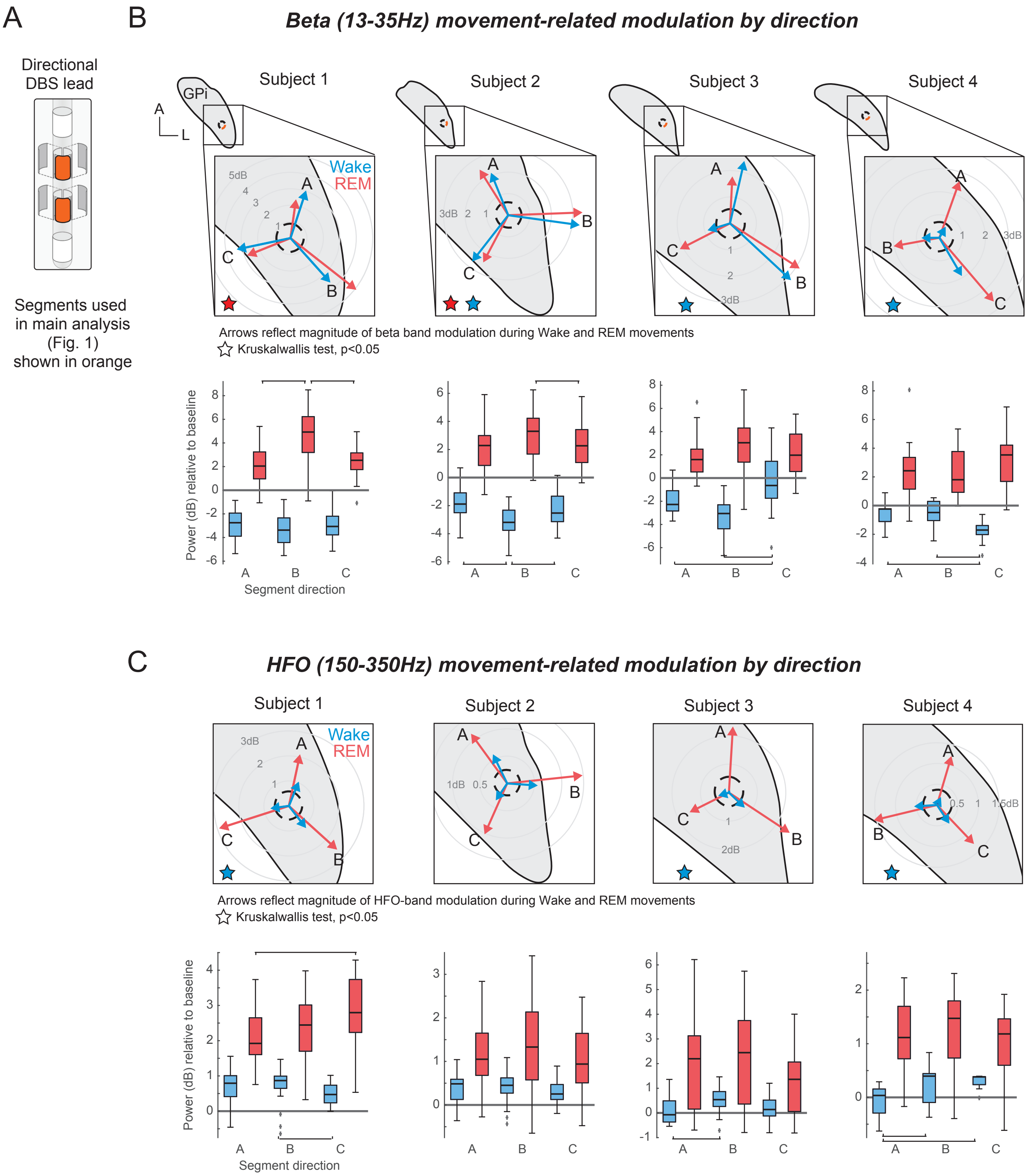

A
B
Beta (13-35Hz) movement-related modulation by direction
Directional DBS lead
Segments used in main analysis (Fig. 1) shown in orange
Subject 1
Subject 2
Subject 3
Subject 4
GPi
Wake
REM
Arrows reflect magnitude of beta band modulation during Wake and REM movements
Kruskalwallis test, p<0.05
Power (dB) relative to baseline
Segment direction
C
HFO (150-350Hz) movement-related modulation by direction
Arrows reflect magnitude of HFO-band modulation during Wake and REM movements
Kruskalwallis test, p<0.05
